# Assessing AF2’s ability to predict structural ensembles of proteins

**DOI:** 10.1101/2024.04.16.589792

**Authors:** Jakob R. Riccabona, Fabian C. Spoendlin, Anna-Lena M. Fischer, Johannes R. Loeffler, Patrick K. Quoika, Timothy P. Jenkins, James A. Ferguson, Eva Smorodina, Andreas H. Laustsen, Victor Greiff, Stefano Forli, Andrew B. Ward, Charlotte M. Deane, Monica L. Fernández-Quintero

## Abstract

Recent breakthroughs in protein structure prediction have enhanced the precision and speed at which protein configurations can be determined, setting new benchmarks for accuracy and efficiency in the field. However, the fundamental mechanisms of biological processes at a molecular level are often connected to conformational changes of proteins. Molecular dynamics (MD) simulations serve as a crucial tool for capturing the conformational space of proteins, providing valuable insights into their structural fluctuations. However, the scope of MD simulations is often limited by the accessible timescales and the computational resources available, posing challenges to comprehensively exploring protein behaviors. Recently emerging approaches have focused on expanding the capability of AlphaFold2 (AF2) to predict conformational substates of protein structures by manipulating the input multiple sequence alignment (MSA). These approaches operate under the assumption that the MSA also contains information about the heterogeneity of protein structures. Here, we benchmark the performance of various workflows that have adapted AF2 for ensemble prediction focusing on the subsampling of the MSA as implemented in ColabFold and compare the obtained structures with ensembles obtained from MD simulations and NMR. As test cases, we chose four proteins namely the bovine pancreatic inhibitor protein (BPTI), thrombin and two antigen binding fragments (antibody Fv and nanobody), for which reliable experimentally validated structural information (X-ray and/or NMR) was available. Thus, we provide an overview of the levels of performance and accessible timescales that can currently be achieved with machine learning (ML) based ensemble generation. In three out of the four test cases, we find structural variations fall within the predicted ensembles. Nevertheless, significant minima of the free energy surfaces remain undetected. This study highlights the possibilities and pitfalls when generating ensembles with AF2 and thus may guide the development of future tools while informing upon the results of currently available applications.

## Introduction

Machine learning (ML) has revolutionized the field of protein structure prediction. AlphaFold2^1^ (AF2) is a deep learning model designed for predicting the 3D structures of individual proteins. Solely based on the amino acid sequence as input, it allows the prediction of the tertiary structure of a protein and provides a per-residue score to assess the accuracy of the predictions.^1–3^ AF2 builds on the premise that the conservation of specific residues across evolutionary paths is associated with inter-residual distances and, consequently, with interactions between conserved residues, such as salt bridges, hydrogen bonds, and other intermolecular contacts.^4,5^ It generates multiple sequence alignments (MSAs) consisting of homologue sequences and identifies the conservation and co-evolution of amino acid residues and sequence patterns, facilitating protein structure prediction. AF2 contains of five different neural networks which differ in training time and MSA size, three predict structures solely based on sequence information (models 1.2.1, 1.2.2 and 1.2.3 in AF2 or 3, 4 and 5 in ColabFold), while two neural networks (models 1.1.1, 1.1.2 in AF2 or 1 and 2 in ColabFold) utilized four template structures in addition to sequence information during training. A detailed summary of the differences between the five models can be found in the SI. In AF2s initial step an MSA is generated, AF2 takes a subset of this MSA containing up to 5120 sequences and passes it to the neural network to predict a protein structure. A critical procedure enhancing the predictions of AF2 is an iterative refinement process called “recycling” – in the default setting, AF2 reuses the networks output structure model as input and reruns the network for three times. This yields five structures for each neural network together with scores that describe the quality of the predicted structure (pLDDT – local accuracy and pTM score – global accuracy).^1^ The pLDDT (predicted local distance difference test) score represents a per-residue confidence score of the model compared to the “true” structure, while the pTM (predicted template modelling score) is obtained from a pairwise error prediction computed as a linear projection from the final pair representation.^1^

Many proteins undergo conformational rearrangements across various timescales, which are pivotal for their functionality. Thus, to characterize the biophysical properties responsible for specific protein functions, there is a need to describe proteins as conformational ensembles in solution, rather than as a single static structural model.^6^ Various computational methods, such as molecular dynamics (MD) simulations, have been developed to explore the conformational space of proteins. Unfortunately, MD simulations can be limited by the accessible timescale of large conformational changes, substantially increasing computing time and costs. Enhanced sampling techniques, such as accelerated Molecular Dynamics (aMD)^7^ have been developed to lower energetic barriers and hence be able to sample more of a protein’s free energy surface. However, most of these enhanced sampling techniques are still computationally expensive and often require previous knowledge of the underlying energy landscape. Therefore, it would be beneficial to be able to swiftly predict multiple conformational states and their respective probabilities at low computational cost.

AF2 was as described above initially designed to predict single, static structures and so does not predict two states for proteins known to have two stable folds^8,9^ or model structural differences originating from missense mutations.^10^ However, various pipelines have recently been proposed to utilize AF2 to predict multiple conformational states.^11–15^ Most of these methods focus on the manipulation of the input MSA. For instance, stochastic subsampling of the generated MSA has been shown to enable AF2 to predict structural ensembles of proteins with apo/holo conformational changes.^11^ With this method, the depth of the input MSA and the number of recycling iterations is adjusted. Another approach, called AF-cluster^12^, clusters the sequence space of the generated input MSA by using a density-based clustering algorithm and uses the cluster centers as input MSA for AF2 prediction. SPEACH_AF^13^ reveals alternative states of proteins by damping coevolutionary signals through *in silico* mutagenesis. The Alternative Contact Enhancement (ACE) approach masks certain coevolutionary signals by creating subfamily MSAs.^15^ Recently, AlphaFlow and ESMFlow have been represented as novel tools for mimicking MD simulation combining AF2 and ESM with flow matching.^16^ These approaches enable the sampling of protein ensembles replicating distributions and properties of MD.

In this study, we investigate four different proteins, for which experimental structural information (X-ray and/or NMR) revealed backbone rearrangements on timescales ranging in the low to high microsecond (µs) regime. Employing the stochastic subsampling approach, our aim was to evaluate how modifications of different MSA subsampling sizes and number of recycles influence the quality of predicted ensembles. Furthermore, we characterized the accessible timescales and free energy barriers between captured conformational rearrangements, as estimated by complementary MD simulations.

## Methods

### Datasets

We chose four distinct proteins, the bovine pancreatic trypsin inhibitor protein (BPTI), a thrombin variant, a single domain camelid antibody fragment (nanobody) and a variable fragment (Fv) of an anti-hemagglutinin antibody based on the availability of experimental structures (X-ray or NMR) and MD data, describing critical conformational rearrangements.

#### BPTI (PDB: 1BHC)

Bovine pancreatic trypsin inhibitor protein (BPTI) is a 58 amino acid residue long monomeric protein that inhibits serine proteases (e.g., Trypsin). BPTI is a well-studied protein and has been subjected to analysis by several X-ray crystallography^17–19^ and NMR experiments^20–23^ as well as MD simulations^24,25^ making it an ideal system to test and benchmark ensemble prediction approaches. NMR and MD studies showed that in the nano-to-millisecond timescale sidechain rearrangements of aromatic residues and a disulfide bond isomerization can occur.^20–22,24^ Furthermore, BPTI exhibits a well-characterized backbone rearrangement caused by this disulfide bond isomerization.^24,26^

#### Protease (PDB: 3LU9, 3BEI)

Human thrombin mutant S195A is a thoroughly analyzed system, since it is a crucial participant in the blood coagulation cascade.^27^ It facilitates the cleavage of fibrinogen, which results in the formation of blood clots.^28,29^ Given its pivotal role in blood coagulation, thrombin is an attractive target for drug development, and its structure has been extensively investigated since the late 1980s^30,31^, followed by extensive MD simulations^32^ to uncover the thermodynamics and kinetics of sidechain and backbone rearrangements which correspond to its active (E) and inactive state (E*).^32,33^

#### Immunoglobulins

1) The heavy (H) chain single domain camelid antibody fragment (nanobody) cAb-H7S (PDB: 4M3K, 4M3J) binds to β-lactamase from *Bacillus licheniformis*.^34^ Both available crystal structures of this nanobody show significant differences mainly in the complementary determining region (CDR, region shaping the binding site [paratope]), namely in the CDR-1 and CDR-3 loop. We performed MD simulations of this nanobody to be able to assess the kinetics between possible different paratope states.

2) The anti-hemagglutinin Fv 17/9 influenza antibody exhibits substantial structural rearrangements, especially at the CDR-H3 loop.^35^ There are three crystal structures of the anti-hemagglutinin antibody Fab 17/9 with and without the hemagglutinin fragment (PDB: 1HIM, 1HIN and 1HIL).^36^ In addition, we thoroughly characterized the Fv by means of MD simulations previously^35^, making it an ideal candidate to test the performance of ensemble prediction tools on antibodies.

### Structure Prediction

In this study, protein structures were predicted using ColabFold^37^, a rapid and efficient implementation of AlphaFold2.^1,37^ MSAs were generated with the MMseqs2^38^ based algorithm implemented in ColabFold. To improve structural heterogeneity, we iteratively modified the deepness of the MSA by modifying the *max_seq* (number of randomly selected sequence clusters from the MSA, the first cluster is the query sequence) and *extra_seq* (number of unclustered sequences additionally passed to the main evoformer stack) parameters (using 16:32, 32:64, 64:128 and 256:512 *max_seq:extra_seq* subsampling) and the amount of recycling rounds (Table showing the tested parameters in SI). For all predictions, we used 40 different seeds. The 200 best ranked structures (according to the pLDDT score for monomers and pTM score for multimers) were used for further analysis. Structure models with a pLDDT score above 70 or a pTM score above 0.7 were classified as sufficient, structure models below these values were checked if they contained unfolded and/or misfolded parts. To mimic real-life applications and avoid bias, we did not include templates for the structure prediction.

### Structural comparison of antibody and nanobody CDRs

The structural similarity of selected CDRs of antibodies 1HIN, 1HIM, and 1HIL and nanobodies 4M3K and 4M3J to all available structures with length-matched CDRs was determined. A reference data set was constructed by exporting all antibody structures from a version of the structural antibody database (SAbDab)^39,40^ time stamped to November 30^th^, 2023. Structures were filtered for those solved by X-ray crystallography at resolution lower than 3.5 Å resulting in 7291 Fabs structures from 4198 PDB files after filtering. Structural similarity of CDR-H3 loops of antibodies 1HIM, 1HIL, and 1HIN was calculated to antibody structures in the reference data set with length matched CDR-H3 of 13 residues, containing both heavy and light chains and having no missing residues in the CDR-H3. Structural similarity of CDR-1 and CDR-3 loops of nanobodies 4M3K and 4M3J was calculated to antibody heavy chain and nanobody structures in the reference set with length matched CDR-1 (8 residues) and CDR-3 (14 residues), respectively, and no missing residues in the corresponding CDR. The C⍺ Root Mean Squared Deviation (RMSD) of the CDR residues after alignment on C⍺ atoms of heavy chain framework residues was calculated with code provided by the SPACE2 method.^41^

### Molecular dynamics simulations

As a reference for the conformational ensembles, which we predicted with the above-described ML methods, we relied on MD simulation data. For most of the investigated systems, we compared the predicted ensemble with previously published data (as listed above). The respective applied simulation protocols can be found in the corresponding publications as referenced above. Since there was no conformational reference data available for the single domain camelid antibody fragment cAb-H7S, we performed MD simulations of this system following a well-established simulation protocol (described by Fernández-Quintero et al^42^). Accordingly, we performed 500 ns of well-tempered Metadynamics simulations using the GROMACS simulation software package together with the PLUMED 2 implementation.^43–48^ As a collective variable (CV), we boosted a linear combination of sine and cosine of the ψ-torsion angles of each CDR-loop, which has been shown to thoroughly enhance the underlying motions of the CDR-loops.^35,42,49,50^ The simulations were performed at 1 bar and 300 K in the isobaric isothermal simulation ensemble (NPT). We used a Gaussian bias height of 10 kJ/mol and a width of 0.3 a.u. Gaussian deposition occurred every 5000 steps and a biasfactor of 10 was used. The resulting trajectory was aligned to the C⍺ positions and subsequently clustered based on the two-dimensional RMSD (2D-RMSD) using the average-linkage hierarchical agglomerative clustering algorithm^51^ in cpptraj^52^ with a RMSD cutoff criterion of 1.5 Å. The obtained 429 cluster representatives were used as starting conformations for 100 ns MD simulations using the AMBER 20 simulation software package which contains the GPU implementation of the Particle Mesh Ewald MD method (pmemd.cuda).^53^ Bonds involving hydrogen atoms were restrained using the SHAKE^54^ algorithm, allowing a timestep of 2.0 femtoseconds. The systems pressure was maintained at 1 bar by applying weak coupling to an external bath using the Berendsen algorithm.^55^ The Langevin Thermostat was utilized to keep the temperature at 300 K during the simulations.^56^ For comparison, we additionally used the AF2 predicted ensembles as starting conformations for 100 ns MD simulations following the same simulation protocol as mentioned above.

### Further analysis

Analysis of the predicted ensemble included a root-mean-square deviation (RMSD) analysis. RMSD calculations as implemented in cpptraj^52^ were performed on C⍺ atoms to compare local or global differences from a reference structure. To compare the obtained trajectories with the predicted ensemble, we performed principal component analysis (PCA) and time-lagged independent component analysis (tICA) with a lag time of 10 ns using the python library PyEMMA 2.^57^ PCA was applied to filter the largest uncorrelated movements observed during the simulation, while tICA was applied to identify the slowest movements of the investigated systems and consequently to obtain a kinetic discretization of the sampled conformational space. Thermodynamics and kinetics were calculated with a Markov-state model (MSM)^58^ with a lag time of 10 ns using the PyEMMA 2 package^57^, which uses the k-means clustering algorithm^59^ to define microstates and the PCCA+ clustering algorithm^60,61^ to coarse grain the microstates to macrostates. To visualize the predicted structures, we used PyMOL (Version 2.4.0).^62^

## Results

In this study, we tested the capability of AF2 with MSA subsampling to predict structural ensembles of various types of proteins using the stochastic subsampling approach. We further analyzed the predicted structures with available experimental and MD data. For all predicted ensembles, pLDDT and pTM scores of each rank can be found in the SI.

### 1. BPTI ensembles predicted with AF2 do not capture all observed conformational states

BPTI is a small (58 amino residues) and globular protein with three characteristic disulfide bonds and well-defined secondary structure. We used NMR data (PDB accession code: 1UUA)^23^, which contains the above-mentioned structural diversity, as reference. We also compared the predicted ensemble with the selected available crystal structures (PDB: 1BHC)^19^. Furthermore, we used the kindly provided 1 ms MD simulation of BPTI performed by the D.E. Shaw research group to compare the predicted AF2 ensembles with the ensembles obtained from MD simulations.^24^ This comparison allows us to determine whether the predicted structures are similar to the most dominant states and if they are at all present in the sampled conformational ensemble from the simulation.

#### 1.1 Structure prediction shows bias towards crystal structures

The ensemble prediction utilizing three recycling rounds yielded structure models similar compared to the crystal structures (PDB: 1BHC) with C⍺-RMSD values ranging from 0-2 Å (Figure 1A). In the case of the shallowest subsampling (16:32), lower-ranked predictions (around rank 125 to 200) exhibit higher RMSD (over 4 Å) compared to the crystal structure (PDB: 1BHC) and lower pLDDT (under 80, Figure SI 2). After visual inspection we found misfolded or unfolded regions for structures with low pLDDT. Local RMSD calculations were performed to evaluate the presence of disulfide bond isomerization between residue 14 and 38 (highlighted in Figure 1C) in the predicted ensemble, along with potential backbone rearrangements near this disulfide bridge, the ensembles were aligned to the respective residues in the crystal structure and local RMSD calculations only considering the respective residues (as listed in the Figure 1 caption) were performed (Figure 1B). In contrast to global RMSD, lower-ranked structures for the 16:32 and 32:64 subsampling display increased RMSD values, indicating a backbone rearrangement of the investigated region. This increased flexibility was not only found for the investigated region, but also for residues 43 to 49 (Figure SI 4). Since the structural heterogeneity for runs with recycling was relatively poor, predictions without recycling runs were also included (Figure 1, labeled as “norec”). Generally, the absence of recycling and a shallow MSA results in larger fluctuation in the backbone region of interest. The presented RMSD plots also indicate that the absence of recycling leads to structures of lower quality, particularly with shallower subsampling of the MSA (Figure SI 2). Among all predictions, the highest-ranked structure models are predicted with the AF2 models 4 and 5 (Figure SI 2). The predicted ensemble of the 32:64 subsampling without recycling can be found in SI (Figure SI 13).

**Figure 1:**
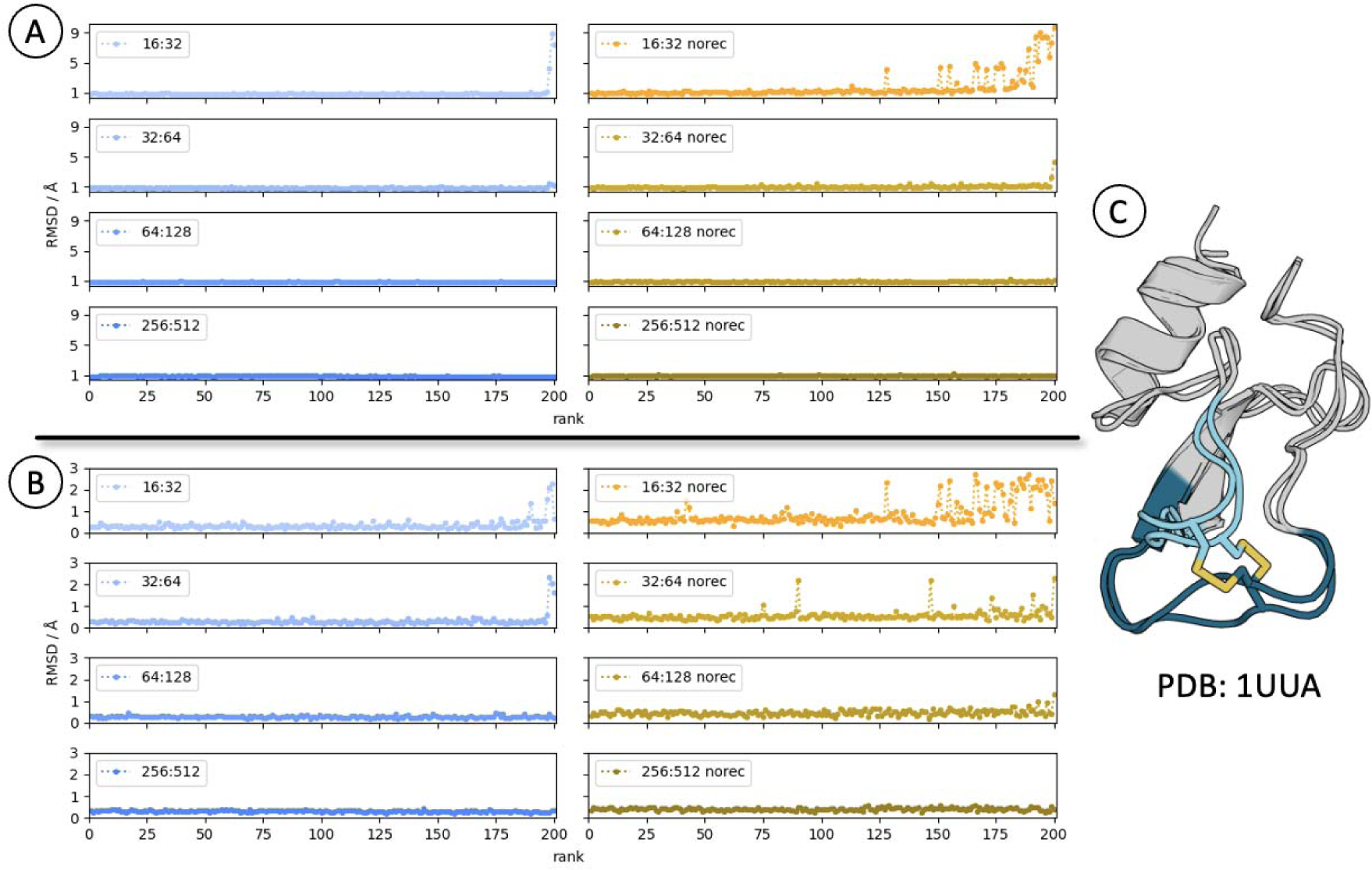
**(A)** Influence of the recycling parameter on the global C⍺-RMSD of the 200 best ranked AF2 predicted structures for BPTI. A crystal structure (PDB accession: 1BHC) was used as reference structure for the RMSD calculations. The blue plots include AF2 ensemble predictions with three recycling rounds, while the yellow plots illustrate RMSD results for AF2 ensemble predictions without recycling rounds. **(B)** Influence of the recycling parameter to the local C⍺-RMSD of residues 10 to 17 (darker blue in Figure 1C), a region that shows backbone rearrangements in the predicted AF2 ensemble as well as in MD simulations and NMR data. The crystal structure (PDB accession: 1BHC) was used as reference structure for the RMSD calculations. Again, the left plots include AF2 ensemble predictions with three recycling rounds, while the right plots illustrate RMSD results for AF2 ensemble predictions without recycling rounds. **(C)** Structural representation of the possible backbone rearrangements when comparing two NMR structures (PDB accession: 1UUA). In darker blue are the residues 10 to 17, whereas in light blue are residues 33 to 40 shown. The disulfide bond affected of isomerization is highlighted in yellow sticks.

#### 1.2 Predicted ensembles miss side minima of the free energy surface occurring in MD

We next performed an analysis of the experimentally determined structures. To determine the position of the available NMR and X-ray structures, we projected the NMR ensemble (PDB accession: 1UUA) and the crystal structure (PDB accession: 1BHC) in both the PCA and the tICA space of selected backbone torsions of the provided MD simulation (Figure 2).

**Figure 2:**
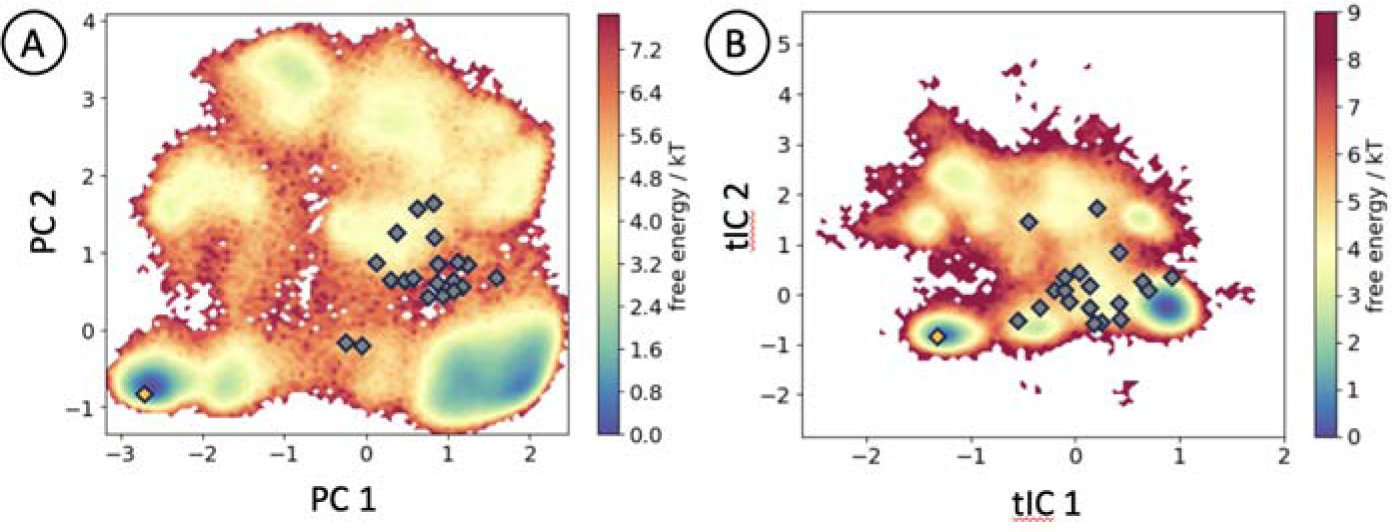
**(A)** PCA and **(B)** tICA analysis of φ and ψ backbone torsions of residues surrounding the 14-38 cysteine disulfide bridge (including residues 10 to 17 and 33 to 40) of BPTI for the 1 ms MD simulation and projection of selected crystal (yellow, PDB: 1BHC) and NMR (grey, PDB: 1UUA) structures of BPTI. The crystal structure lies in a dominant minimum of both PCA and tICA space. The NMR structures cover a different conformational space in both PCA and tICA, for (A), the NMR structures are not located near or in a dominant minimum, while for (B) the NMR structures show a distribution alongside tIC1.

We found that the NMR structures occupy a distinct conformational space in the backbone torsions of residues surrounding the 14-38 cysteine disulfide bridge when compared to the crystal structure. In the PCA plot (Figure 2A), the NMR structures do not lie within the dominant minimum sampled by the crystal structure. Additionally, a second dominant minimum, observed alongside PC1 and tIC1, disagrees with structural data. Below, the ML-generated ensembles with the MD simulation are compared analogously. Thus, to evaluate whether the predicted ensemble conforms to the free energy surface of the region of interest, we performed a PCA and tICA analysis and projected the ensembles onto the respective surface. We compared the AF2 predicted ensemble again with the 1 ms long MD simulation conducted by the D.E. Shaw research group.^24^

Figure 3 shows that structure models generated with three recycling rounds (blue dots) are predominantly located within a dominant minimum in both the tICA and PCA plots. AF2 predictions fail to capture alternative structure models located within a dominant minimum alongside PC1 (representing biggest movement) and tIC1 (representing slowest movement). Overall, the utilization of recycling and deeper MSAs increases the preference towards crystal structures.

**Figure 3:**
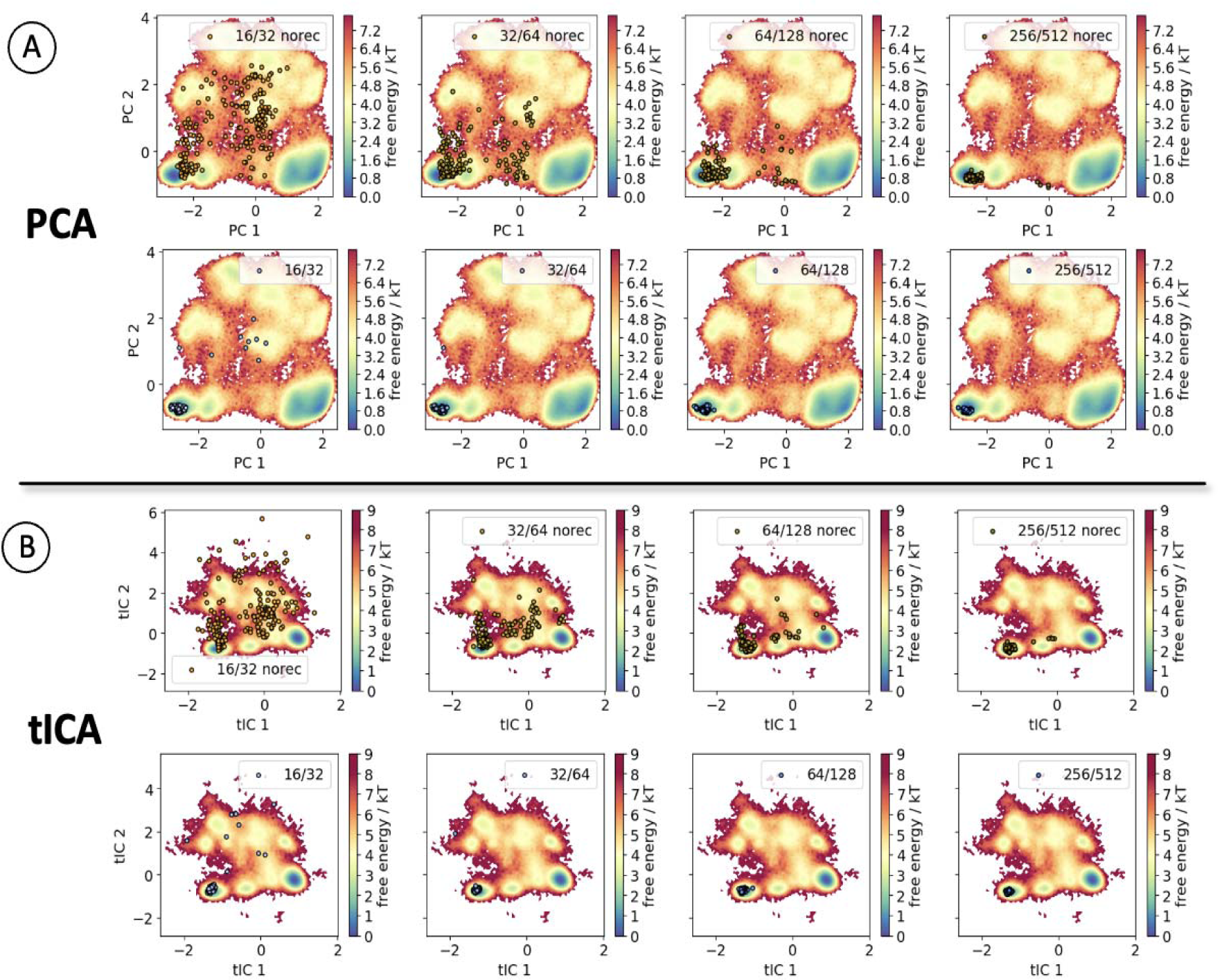
PCA and tICA analysis of φ and ψ backbone torsions of residues 10 to 17 and 33 to 40 of BPTI for the 1 ms MD simulation, including the projected AF2 predicted ensembles into the respective space as yellow (for the predictions without the recycling runs) or blue (with recycling runs) dots. **(A)** shows the PCA space, while **(B)** shows the tICA space.

**Figure 4:**
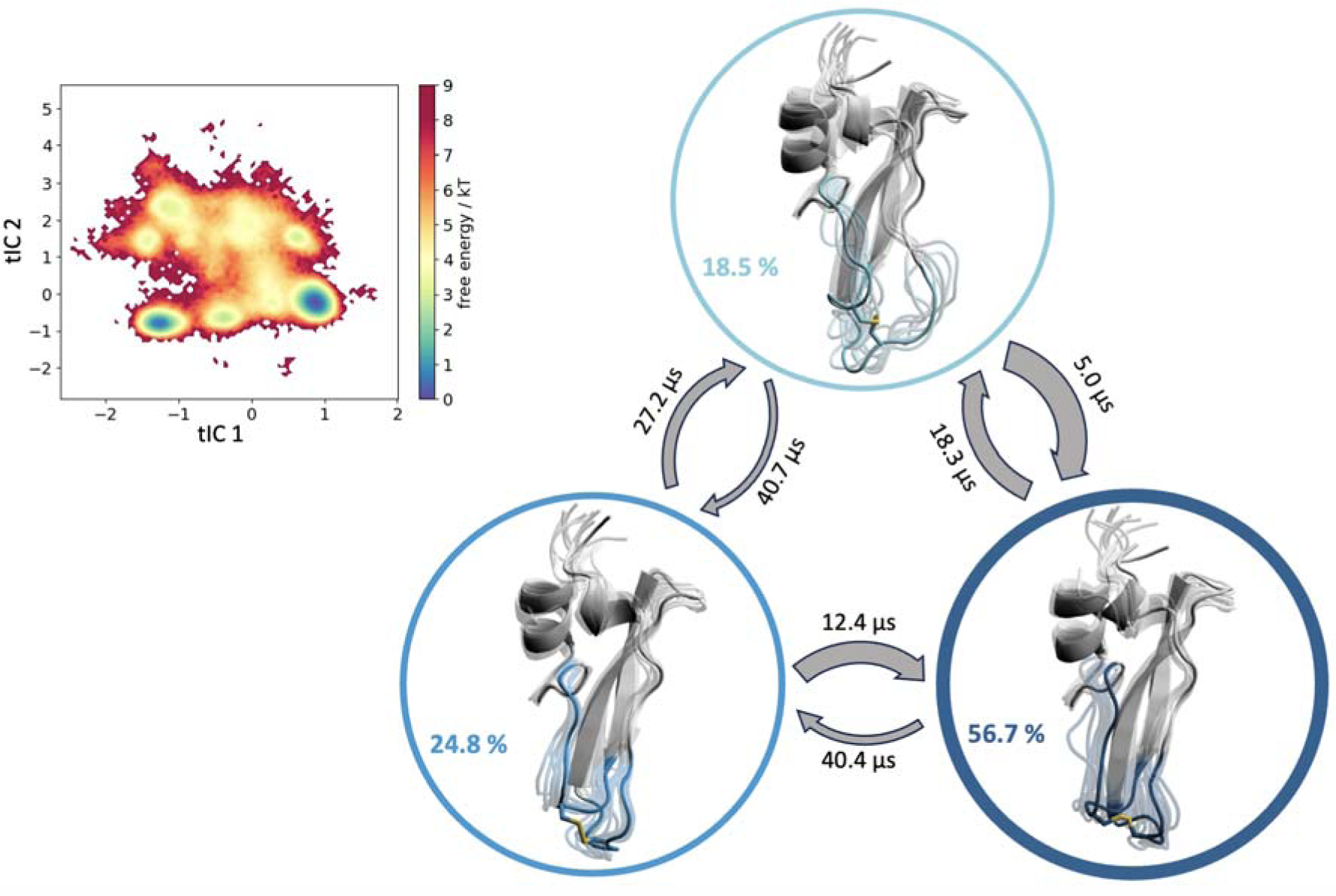
Reweighted tICA analysis of φ and ψ backbone torsions of residues 10 to 17 and 33 to 40 of BPTI for the 1 ms MD simulation as well as the Markov-state model (MSM) to estimate thermodynamics and kinetics of the respective conformational rearrangements, resulting in three different macrostates. The probabilities of each state are given in percent. The highest populated state (colored in dark blue) represents the alternative backbone conformation around the switched disulfide bond (yellow). The transition times between the states are on the µs timescale.

Disabling the recycling results in structure predictions that not only align within the same global minimum as the crystal structures but also remain on the overall free energy surface. Some of the lower-ranked structures, especially for the 16:32 subsampling, deviate from the estimated tICA free energy surface, as indicated by the RMSD analysis, suggesting potential misfolded regions. Notably, structures along PC1 and tIC1 are predicted for all subsamplings, yet the above-mentioned dominant minimum remains uncaptured (Figure SI 3). Ensembles without recycling contain structures covering a similar conformational space as the NMR structures in both PCA and tICA.

To discretize the conformational space and to quantify the timescale of rearrangements captured in the 1 ms MD simulation of BPTI, we performed Markov-state models (MSM) and find transitions in the low microsecond timescale for the characteristic motions of BPTI known from NMR and X-ray structures. One of the prominent movements of BPTI, namely the backbone rearrangements which follow the isomerization of the disulfide bridge, is represented by the tIC1, and is not sampled within the predicted AF2 ensemble.

### 2. Human thrombin mutant S195A ensemble predictions do not sample the inactive state

Thrombin plays a pivotal enzymatic role in plasmatic blood coagulation. Its activity is regulated by Na^+^-ions, with its binding site situated within the Na^+^-loop.^63^ Structural heterogeneity between active and inactive thrombin is primarily observed in the region of the Na^+^-loop, which includes residues 213 to 229.^64^ The timescales of transitioning between the active and inactive form can vary significantly (over tenfold) and are dependent on the presence of Na^+^ ions.^32^ With numerous crystal structures available for both the active and inactive form (PDB: 3LU9 as representative for the active form and 3BEI for the inactive form, all other PDBs are provided in Table SI 3), alongside with available crystal structures of thrombin in its zymogen form, AF2 can be used to predict a structural ensemble that spans the free energy surface of this protein (Figure 5B).

**Figure 5:**
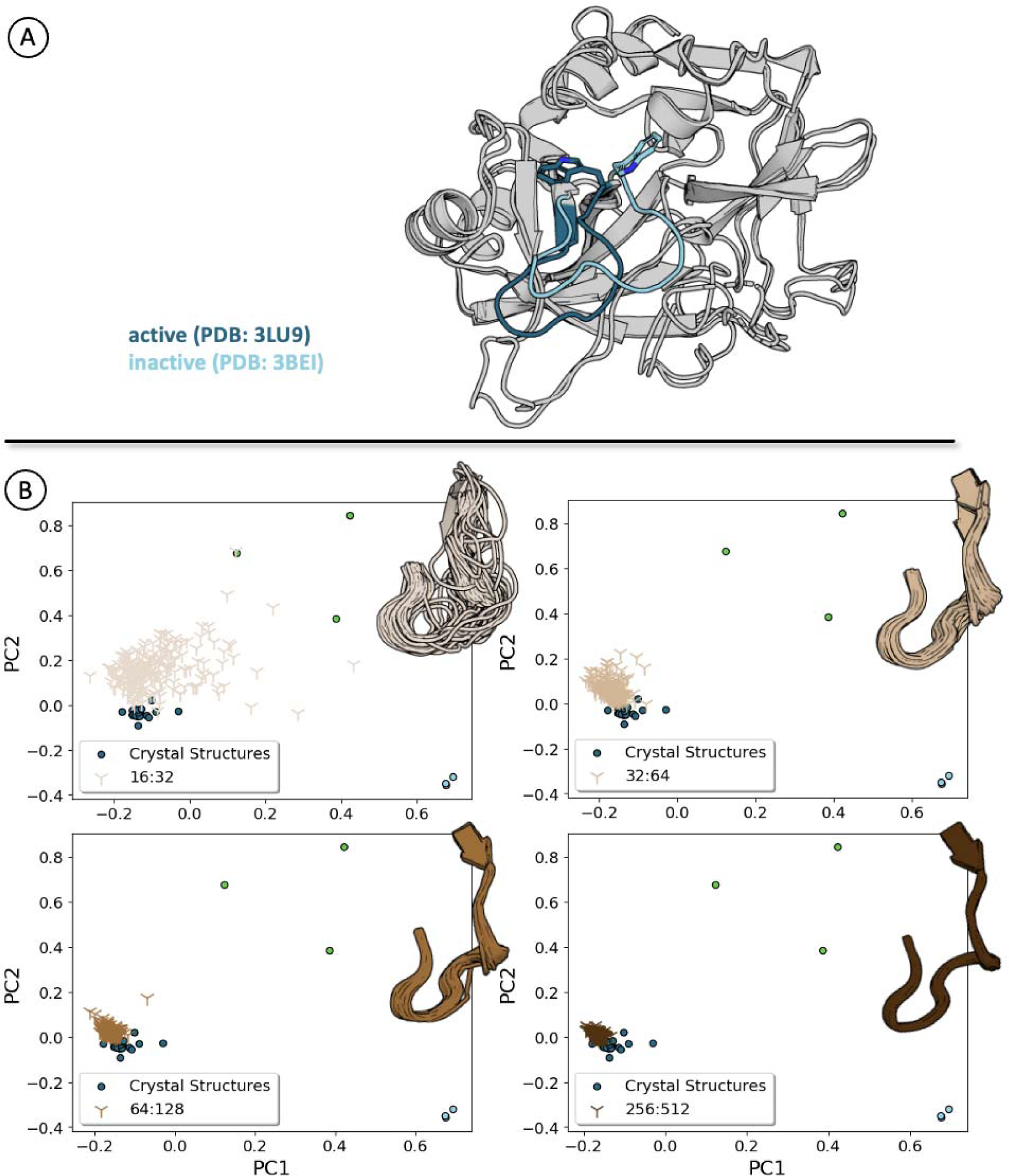
**(A)** Structural representation of the active (E) and inactive (E*) form of thrombin. Structural differences can be seen for the Na^+^-loop, which is colored in dark blue (active) and light blue (inactive). The reorientation of W215 is also depicted as sticks. **(B)** PCA analysis of the available crystal structures (used PDB entries are listed in Table SI 3) of thrombin and projection of AF2 generated ensembles using varying subsampled MSAs (the deeper the MSA, the darker the color). As feature, the C⍺ positions of residues 212 to 229 were chosen. Structures were aligned to the active form (PDB: 3LU9). Crystal structures representing the active conformation are colored in dark blue, while the inactive form is colored in light blue. Zymogen proteins are colored in green. Next to every PCA plot, a structural representation of the Na^+^-loop of the AF2 generated ensemble is shown.

#### 2.1 Structural diversity in the predicted ensemble is missing when shallowing the MSA

We utilized the stochastic subsampling approach with diverse parameters to evaluate the capacity of AF2 to predict active and inactive structures independent of any templates. Subsequently, we performed a PCA analysis of the available crystal structures and projected the AF2 predicted ensemble onto the PCA space.

Even with the shallowest subsampling examined, the overall conformation of the loop region fails to resemble the inactive form. Residue W215, which undergoes a sidechain flip between active and inactive form (Figure 5A), demonstrates increased flexibility with shallower subsampling. While other loop regions of thrombin exhibit a high variance in possible conformations with lower subsampling, the Na^+^-loop shows minimal variance. Lower-ranked predictions for the shallowest subsampling exhibit low pLDDT and pTM scores, which should be carefully checked for physical inaccuracies (Figure SI 6). The highest ranks are predicted with the AF2 model 5, one of the template-refined models.

### 3. Nanobody CDR loop state transitions with AF2 between bound and unbound crystal structures

Camelid heavy chain-only antibody variable domains, popularly known as nanobodies, are one of the smallest-known functional antibody fragments with vast applications within both therapy, diagnostics, food supplements, and feed additives.^65,66^ Capturing nanobody movements is crucial, particularly considering the CDR loops, which are largely responsible for antigen binding. For nanobodies, the CDR-3 loop is especially important for the binding interactions and is the most diverse loop. This specificity of antigen recognition has been linked with CDR-3 flexibility and to its longer chain length compared to the equivalent CDR loops of conventional antibodies. Moreover, its high flexibility also allows this loop to bind cryptic epitopes buried within the antigen.^42,67,68^

Predicting nanobody structures using AF2 can be challenging due to the high sequence similarity of nanobodies to conventional antibody heavy-chain variable domains, as AF2 utilizes originally paired heavy-chain variable domains from Fvs as templates. This similarity can potentially influence the predicted shape of the CDR loops and ultimately lead to artifacts in the prediction.^69^ Furthermore, immunoglobulins are a highly conserved class of proteins, and mutations introduced in the CDR loops often diverge from evolutionary paths. Therefore, ensemble predictions for nanobodies may not benefit from the recently developed AF2 ensemble predictions tools that focus on the modification of the MSA. As an example, we here present a nanobody (single domain camelid antibody fragment cAb-H7S which binds to β-lactamase from *Bacillus licheniformis*) with two X-ray structures available (PDB: 4M3K and 4M3J) (Figure 6B), which differ in the conformations of their CDR-1 and CDR-3 loops.

**Figure 6:**
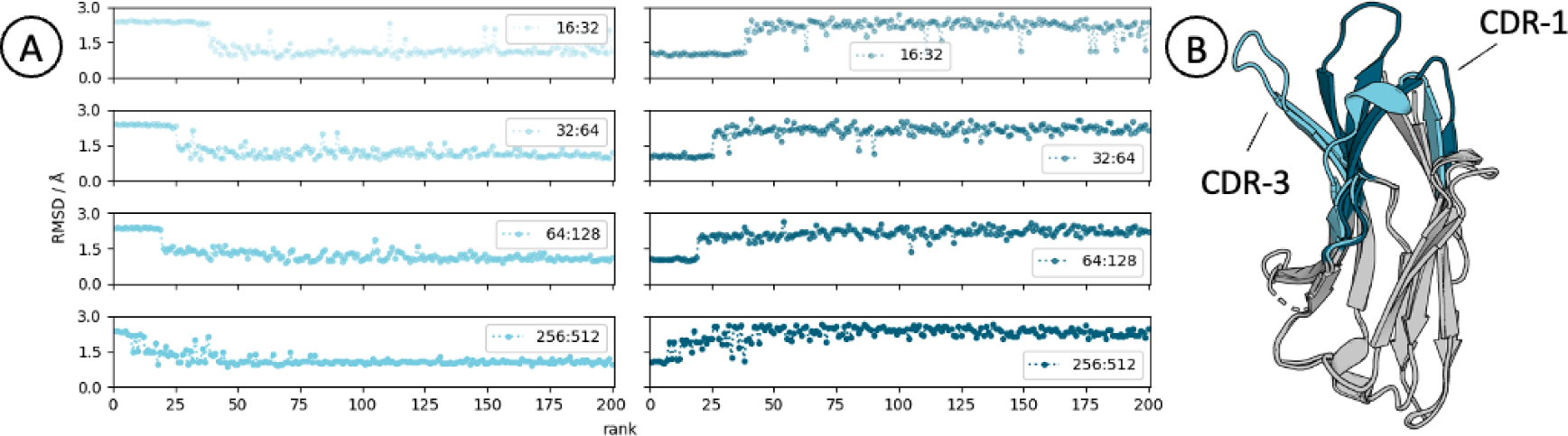
**(A)** RMSD calculation of all C⍺ atoms. The ensemble structures were aligned to the unbound (PDB: 4M3J, colored in light blue) and bound (PDB: 4M3K, colored in dark blue) crystal structures. **(B)** The two crystal structures are depicted, and the CDR loops are colored in the corresponding color (CDR-1 in dark and CDR-3 in light blue).

#### 3.1 MSA deepness influences ratio of bound/unbound AF2 ensemble predictions

Examining the RMSD plot of all C⍺ atoms, it becomes apparent that the top ranked structures resemble a conformation similar compared to the crystal structure of the nanobody in complex with the target (PDB: 4M3K), whereas lower ranks show a similar structure to the nanobody free in solution (PDB: 4M3J). Remarkably, reducing the depth of subsampling yields an increased prediction of models more similar to the bound structures. The predicted ensemble of the 32:64 subsampling can be found in SI (Figure SI 13).

#### 3.2 Comparison with MD data uncovers sampling of certain CDR-1 dynamics with AF2 ensembles

We evaluated whether the predicted ensembles are situated within the free energy landscape of the nanobody and analyzed the effectiveness of AF2 utilizing MSA subsampling in predicting diverse conformations of the CDR loops. Our assessment entailed a comparison of the conformational landscape observed during the MD simulation, specifically focusing on tIC1 and tIC2 components, with the ensemble forecasted by AF2.

Figure 7 depicts the PCA and tICA based on the backbone torsions for the CDR-1 and the CDR-3 loop with the crystal structures projected into the respective space. Due to the backbone torsions used as feature, the bound and unbound crystal structure lie in different minima in both tICA and PCA for CDR-1, while the CDR-3 structures fall into the same minimum. Whilst all ensemble models align within a dominant minimum for the CDR-3 loop, greater variance is observed for CDR-1. CDR-1 comprises of several side minima, with the ensemble prediction covering some of them, while the bound and unbound crystal structure reside in different minima. The pLDDT and pTM scores are deemed sufficient for all subsamplings (Figure SI 6).

**Figure 7:**
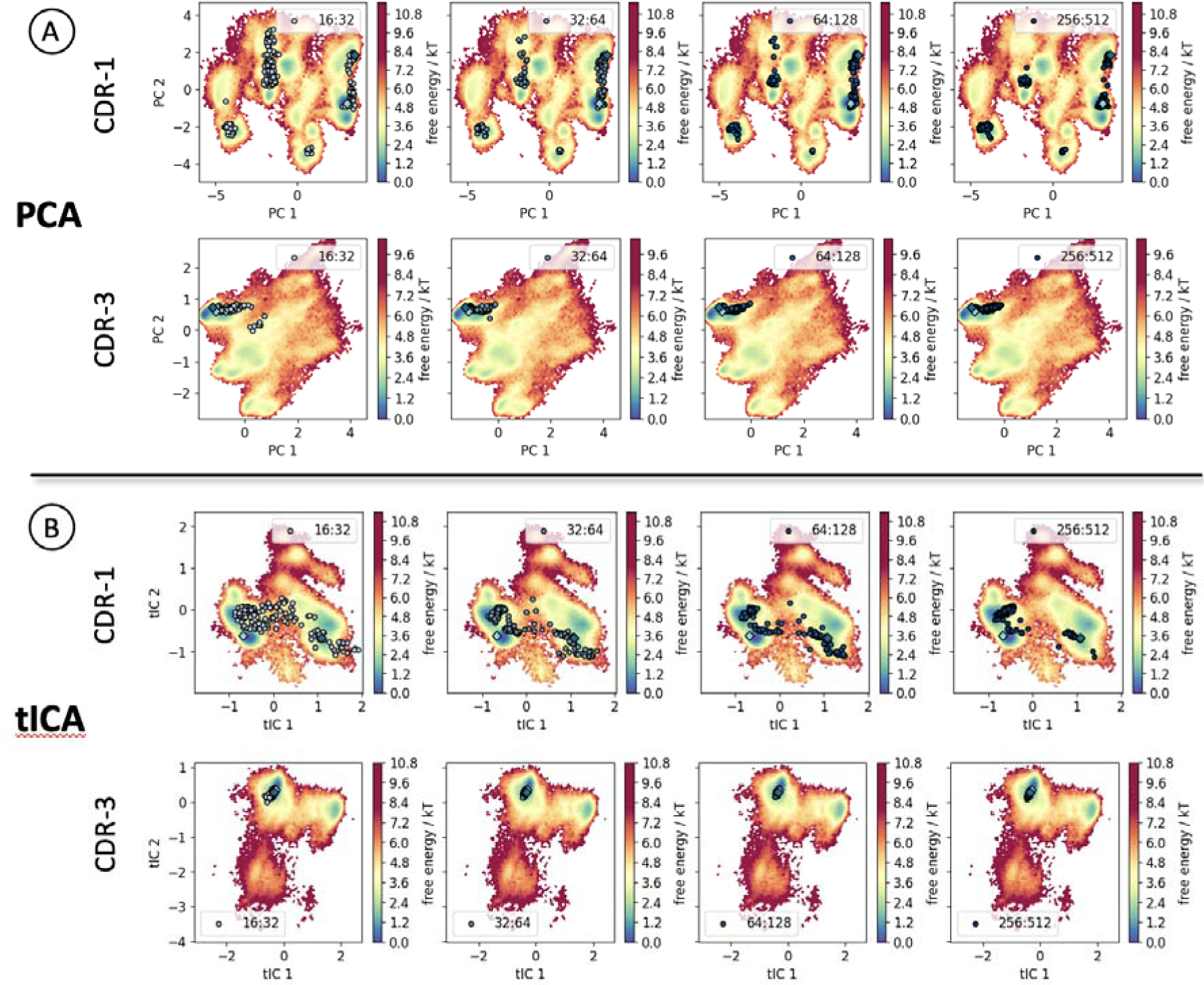
Comparison of the predicted AF2 ensembles with MD data. **(A)** PCA and **(B)** tICA are performed based on the φ and ψ backbone torsions of the CDR-1 (top) and CDR-3 loop (bottom), respectively, and the AF2 predicted ensemble (depicted as circles) is projected onto the respective PCA and tICA space. With increasing depth of the MSA the colors of the projected predicted AF2 ensemble get darker. The bound (dark blue diamond) and unbound (light blue diamond) crystal structures lie in minima.

To reconstruct the kinetics of the CDR-1 loop conformational change upon binding, an MSM was conducted, as illustrated in Figure 8, which depicts the corresponding transition times.

**Figure 8:**
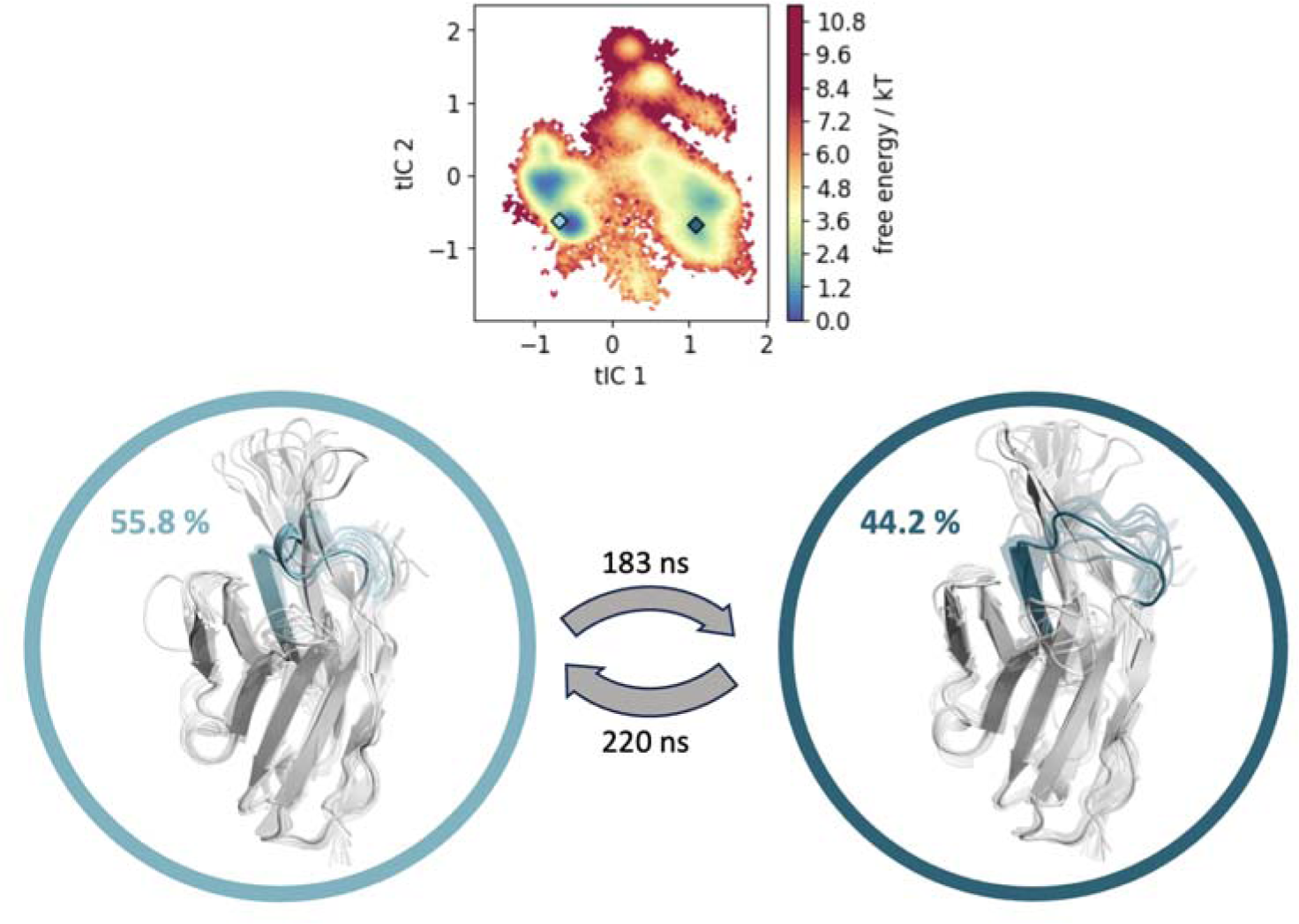
MSM based on the φ and ψ backbone torsions of the CDR-1 loop to estimate state probabilities and transition kinetics of the respective conformational rearrangements of the cAb-H7S nanobody. The MSM results in two macrostates, the highest populated state corresponds to the unbound conformation (PDB: 4M3J, colored in light blue), the second state shows a conformation similar to the bound crystal structure (PDB: 4M3K, colored in dark blue). The resulting transition times between the macrostates occur on the high nanosecond timescale.

### 4. AF2 captures the unbound but not the binding competent conformation of the CDR-H3 loop

As for nanobodies, antibodies pose challenges in protein structure prediction.^70^ While five of their six CDR loops tend to adopt different canonical conformations, CDR-H3 has proven to be difficult to model due to its increased diversity in sequence and length.^71^ Here, we examine an antibody (anti-hemagglutinin Fv 17/9 influenza antibody) that exhibits distinct conformations in the CDR-H3 (Figure 9B). We use previously published MD data^35^ to compare the AF2 predicted ensembles with the free energy surface of the CDR-H3. Additionally, we compare the predicted ensembles with three available crystal structures (PDB accession: 1HIM/1HIN/1HIL), representing both bound and unbound conformations.^36^

**Figure 9:**
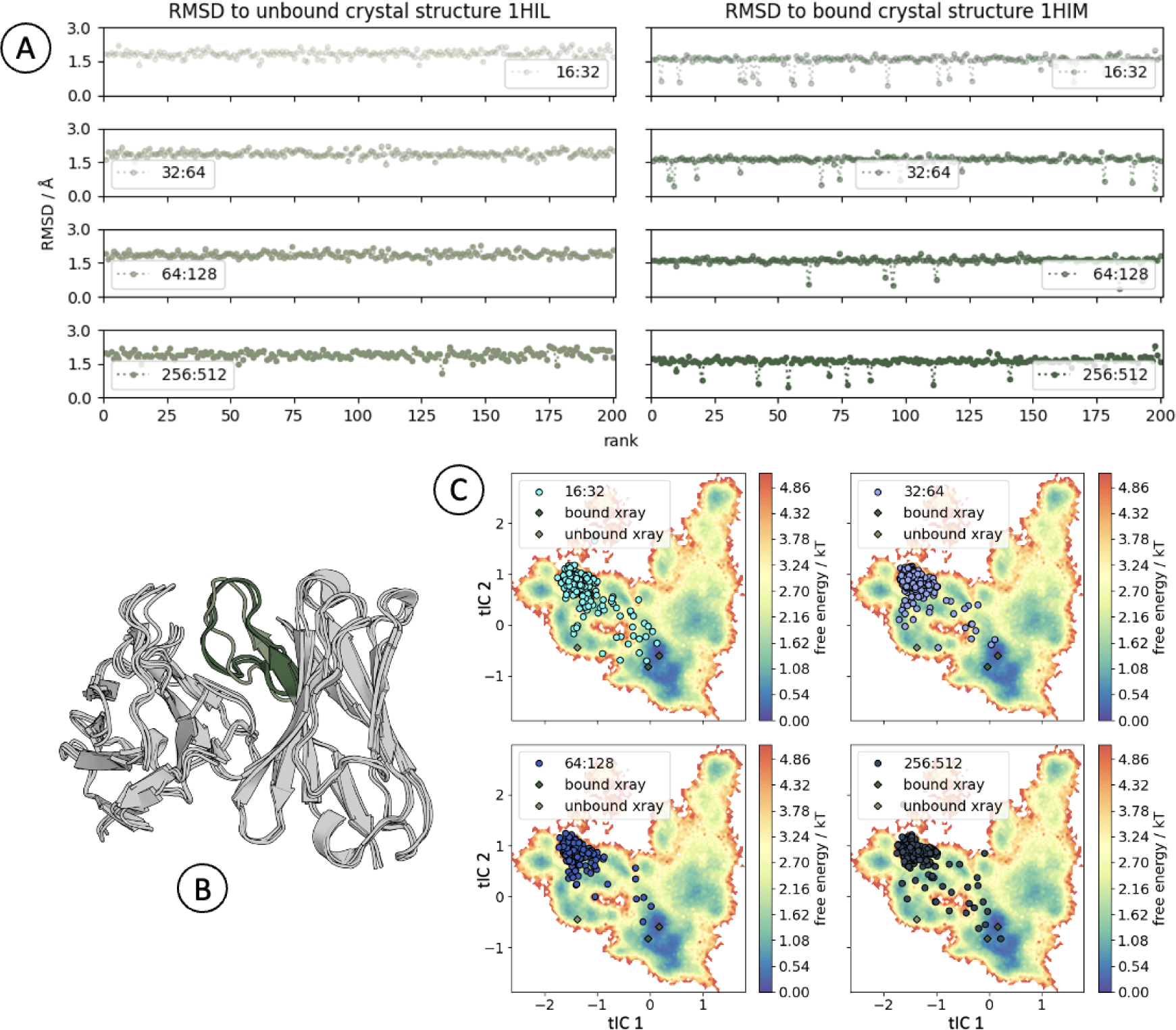
**(A)** CDR-H3 loop (Chothia nomenclature) C⍺ RMSD analysis of the predicted ensemble in reference to the unbound (light green, PDB accession: 1HIL) and bound crystal structure (dark green, PDB accession: 1HIM, 1HIN). **(B)** Overlay of the three available crystal structures of the Influenza Hemagglutinin Fv. In dark green, the CDR-H3 loops in the bound state are depicted, while the unbound conformation is colored in green. **(C)** tICA analysis based on the aligned C⍺ positions (reference: bound crystal structure, PDB: 1HIM) of the CDR-H3 including the projection of the crystal structures (diamonds) and the AF2 predicted ensembles (dots).

#### 4.1 Vast majority of CDR-H3 conformation predictions differ from structural data

Figure 9A illustrates local RMSD calculations of CDR-H3, which show that individual structures are predicted in the bound conformation, while most of the AF2 predicted ensemble represents a state, which differs from both the unbound and bound crystal structure. In this case, the deepness of the MSA does not affect the number of structures predicted in or near the bound conformation.

#### 4.2 AF2 ensemble predictions miss CDR-H3 transitions between bound and unbound conformations

Figure 9C depicts a tICA analysis of the used MD data alongside with the predicted ensembles. The tICA analysis reveals that the majority of the AF2 predicted structures represent a single state of the CDR-H3 across all four subsampling variations. The deepness of the used MSA does not appear to influence the predicted conformational space. More structures are predicted near the unbound state, with the bound state and several side minima distributed along tIC2 barely being captured. The obtained overall pLDDT and pTM values for each predicted structure are considered sufficient (Figure SI 10). The resulting ensemble of the 32:64 subsampling can be found in SI (Figure SI 13). To characterize the timescale of the conformational change between bound and unbound state of the Fv (Figure 9B), we constructed a MSM resulting in transition timescales between these two states in the high microsecond (µs) timescale.^35^

## Discussion

Undoubtedly, AF2 has revolutionized protein structure prediction and significantly enhanced drug discovery and development processes.^72^ Recently, it has been stated that the capability of AF2 to predict structural ensembles can be facilitated by manipulating the multiple sequence alignment.^73^ We assessed this capability across four types of proteins and subsequently compared these ensembles with available experimental and/or MD data to determine a correlation between the ensembles and the underlying free energy surface of the respective MD simulation. Depending on the type of protein and the timescale of a rearrangement, protein dynamics play a pivotal role in determining protein function.^74^ Rapid transitions occurring on the sub-nanosecond timescale exhibit local, small amplitude fluctuations. The energy barrier separating these numerous states is less than 1 kT, contributing to the systems entropy. Conversely, slow transitions describe fluctuations between a low number of states on the µs timescale or slower, with barrier heights of typically several kT. These fluctuations can include loop or bigger domain rearrangements. Slow, kinetically stable states are crucial for many biological processes, including enzyme catalysis, molecular recognition (like protein-protein interactions), and signal transduction.^75^ Especially for antibodies/nanobodies, conformational heterogeneity is a strong determinant of their binding affinity. It has been shown that the binding entropy is an important contributor to the stability of antigen-antibody/nanobody complexes^65,76^ and that characterizing these states can be facilitated through MD simulations.^77^ However, the applicability of MD simulations is constrained by accessible timescales and computational resource limitations. Without observing multiple reciprocal transitions among potential slow conformational changes, assigning Boltzmann weights to these states becomes challenging. Several enhanced sampling techniques can explore different slow states within the configurational space. Nevertheless, successful implementation of these techniques requires prior knowledge of the system to ensure a proper representation of the configurational space of the system.

AF2 subsampling can be used to inform enhanced sampling techniques to optimize parameters for adequate sampling, to circumvent the above-mentioned challenges. Several tools and applications have been published that attempt to tackle these challenges, underscoring the growing importance of ML in unraveling the dynamics of proteins.^78–85^ However, we find here for the considered proteins that AF2 ensembles do not reproduce the free energy surface, and the predicted ensembles can miss important side minima of the conformational space observed in MD. In most cases, more structures are predicted near or within minima.^86^

For BPTI, we found that there is a noticeable bias towards crystal structures – most probably because a crystal structure of BPTI was part of AF2’s training set (release date of a structure of BPTI was before AF2 training cut [2018-04-30]), implying that AF2 predictions may have a bias towards the crystal structure conformations observed in the training set. This has also been found for other proteins.^13^ The highest ranks (based on pLDDT) are consistently predicted by model 5 and the recycling parameter shifts the prediction towards higher structural similarity to the crystal structures. Without recycling, more deviation can be seen, and with shallow subsampling, more structures with low quality (low pLDDT and low pTM) are predicted. Since the pLDDT measures the quality of a structure prediction based on PDB structures, a low pLDDT value does not necessarily evidence a poor prediction, but it might indicate lack of secondary structure elements. Furthermore, the question remains if the recycling procedure removes some more variable coevolutionary signal resulting from the MSA, since the purpose of the recycling procedure is to filter uncertainties from each model and refine regions that the network is less confident about. Comparing the transition times from the Markov-state model with the obtained ensembles, AF2 can predict ensembles that lie within the nanosecond and low microsecond timescale, however, it does not capture the well-characterized isomerization of the disulfide bridge for BPTI. Even after starting MD simulations from the 200 conformations obtained by MSA subsampling, we do not capture the most dominant minimum, corresponding also to states in NMR, from the 1 ms reference MD simulation provided by the D.E. Shaw research group (Figure SI 3).

For thrombin, we find predictions for the active (E) state. However, AF2 fails to predict the inactive (E*) state. This is somewhat surprising, since various crystal structures of bound and unbound thrombin are available. It might be that just the bound state of thrombin was in the training set of AF2. Previous studies characterized the conformational transition from E-like to E*-like states with MD simulations.^32^ The equilibrium between the active and inactive forms of thrombin have been discussed to be governed by the allosteric binding of a sodium ion. Long time-scale MD simulations showed a sodium-induced conformational shift, resulting in a stabilization of the active form in presence of a Na^+^-ion and a shift towards the inactive form in sodium-free simulations.^32^ Predicted structures with AF2 lack coordinates of solvent molecules, ions, cofactors, ligands and do not take post-translational modifications into account. ^87^ However, AF2 is still able to accurately predict the E state, which has been described to be stabilized by the functionally critical Na^+^-ion, while not capturing the inactive state which would be favored, without the presence of the Na^+^-ion. One potential explanation could be that within the PDB the number of active state structures is higher, biasing the MSA subsampling towards the activated state even without a Na^+^-ion. The top ranked structures (based on pLDDT) are all predicted with model 4 and model 5. This trend is similar to that seen for BPTI ensemble prediction, which questions whether model 4 and model 5 are suited for ensemble predictions (see Table SI 2).

In the case of immunoglobulin fragments, the depth of subsampling does not appear to significantly influence the conformational space covered by the ensembles. We hypothesize that this is caused by the missing coevolutionary signal in the CDR loops of immunoglobulins.

Regarding the nanobody, while the highest ranks are overall in proximity to the bound crystal structure, local analysis reveals that the highest ranks show CDR-1 structures similar to the unbound crystal structure conformation (PDB: 4M3J). This behavior also manifests in structure models originating from the NanoBodyBuilder2 model, a nanobody-specific model (Figure SI 9).^88^ Subsampling appears to affect the ratio of predicted unbound to bound structures. The most prominent disparity in the ensembles is observed in the CDR-1 loop (Figure SI 7 and SI 8). This conformational rearrangement can also be seen in the available crystal structures. Figure SI 7 shows the comparison of the conformational space of the CDR-1 loop obtained from our in-house MD data with the space covered by starting MD simulations from the 200 predicted structures from AF2. While the CDR-1 conformational space starting from the 200 predicted AF2 structures is smaller compared to the in-house MD, the highest populated states along tIC1, corresponding to the conformational changes observed in the available X-ray structures, are captured with comparable probabilities.

Similar challenges in capturing relevant conformational changes in the CDR loops as for nanobodies also apply for antigen-binding fragments. Figure 9 displays the comparison of the predicted AF2 ensemble with 11 µs of classical MD simulations and shows that MSA subsampling of AF2 does not succeed to capture the bound conformation of the Fab 17/9. This contrasts with results obtained from the ABodyBuilder2 model, which was specifically developed to predict antibody structures (Figure SI 12).^88^ Also starting MD simulations from the predicted AF2 ensembles did not further improve the sampling towards the bound conformation (Figure SI 11). Figure SI 11 (bottom) shows the projection of structurally similar structures present in the PDB to the available X-ray structures of the CDR-H3 loop of the Fab 17/9. The fact, that no PDB structures with a similar structure to the bound CDR-H3 loop conformation could be found, can be one determinant why AF2 in combination with MSA subsampling did not capture this conformational transition. This agrees with the findings for the nanobody, where we actually capture the experimentally observed structural changes upon binding with the predicted AF2 ensemble for the CDR-1 loop and show that similar structures to both the bound and unbound conformational states are present in the PDB (Figure SI 7), emphasizing the critical role of available structural data in enhancing the predictivity of ensemble predictions.

In addition to the discussed manipulation strategies of AlphaFold2 MSAs, generative machine learning models have been developed for the task of protein conformation prediction directly.^16,89–91^ These models are trained to learn the distribution of ‘valid’ protein structures which can be sampled to obtain conformational states. Models have mostly been evaluated on case studies and successfully reproduced conformational states for some of them. Among these methods, AlphaFlow^16^ was evaluated on the largest test set containing 100 proteins exhibiting conformational changes. When trained on crystal structure data only, AlphaFlow does not cover more of the conformational space than default AlphaFold2. However, performance is improved when supplemented with additional MD data, offering a promising strategy for conformation prediction when more data on conformational states becomes available.

## Conclusion

In conclusion, regardless of the method employed for ensemble prediction, it is crucial to keep in mind that the quality of ensembles heavily depends on the structural information provided to AF2. This highlights the critical role of accurate structural data in guiding and enhancing the predictive capabilities of ensemble modelling methods in protein structure prediction. Thus, collecting more high-quality structural data is still an important step to advance the field of molecular biology. While subsampling of the MSA and various other methods provide a fast way to generate ensembles of structures, they still do not capture some of the experimentally well-characterized and functionally relevant conformational states for the proteins presented in this study. Even when using the predicted structural ensembles as starting structures for MD, the structural information provided to AF2, e.g., structures in the training set, strongly biases the predictivity and consequently also the sampled distribution and the respective probabilities.

## Funding and Acknowledgments

We acknowledge EuroHPC Joint Undertaking for awarding us access to MeluXina, Luxemburg. The Leona M. and Harry B. Helmsley Charitable Trust (#2019PG-T1D011, to VG), UiO World-Leading Research Community (to VG), UiO: LifeScience Convergence Environment Immunolingo (to VG), EU Horizon 2020 iReceptorplus (#825821) (to VG), a Norwegian Cancer Society Grant (#215817, to VG), Research Council of Norway projects (#300740, #331890 to VG), a Research Council of Norway IKTPLUSS project (#311341, to VG). Funded by the European Union (ERC, AB-AG-INTERACT, 101125630). A.H.L. is supported by a grant from the European Research Council (ERC) under the European Union’s Horizon 2020 research and innovation program [850974] and by a grant from the Villum Foundation [00025302]. A.B.W. was supported by the Bill & Melinda Gates Foundation INV-004923. F.S. is supported by the Engineering and Physical Sciences Research Council (EPSRC), under grant number EP/S024093/1, Roche and the Royal Commission for the Exhibition of 1851.

