## Supplementary_Information for "Assessing AF2’s ability to predict structural ensembles of proteins"

*Table 1: System overview together with tested subsampling modifications of AF2. Further information and explanations about the subsampling and recycling can be found in the introduction of the main text. BPTI stands for* Bovine pancreatic trypsin inhibitor protein, while Fv is the variable fragment of an antibody (Fv 17/9 influenza antibody in this case).

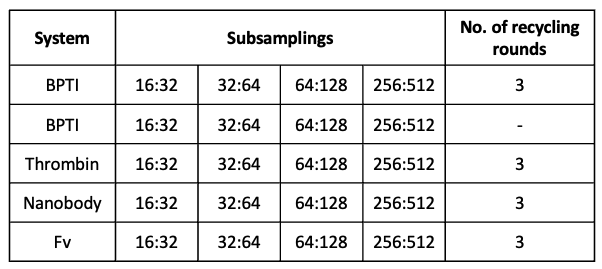

*Table 2: Difference of the five Neural Networks used in AF2 and ColabFold. The main discrepancies lie in the initial form, the number of templates (N. Templates), the max_seq* (number of randomly selected sequence clusters from the MSA, the first cluster is the query sequence) and *extra_seq* (number of unclustered sequences additionally passed to the main evoformer stack)*, the training samples and the training time. Model 2 is a retrained model which took model 1 as initial form, model 4 and 5 are retrained models based on model 3.*

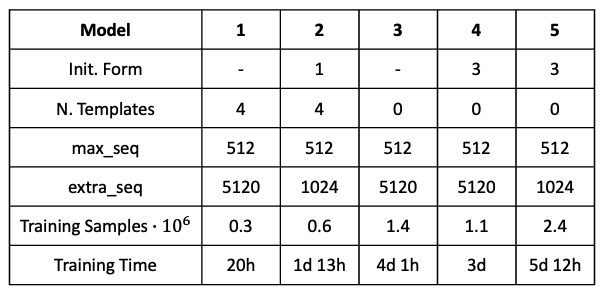

*Figure 1: Evaluation of the AF2 ensemble prediction of BPTI with three recycling rounds. On the left are the pLDDT and pTM score based on the rank depicted, whereas the right plots show the ranks of the different models.*

*Figure 2: Evaluation of the AF2 ensemble prediction of BPTI without recycling rounds. On the left are the pLDDT and pTM score based on the rank depicted, whereas the right plots show the ranks of the different models.*

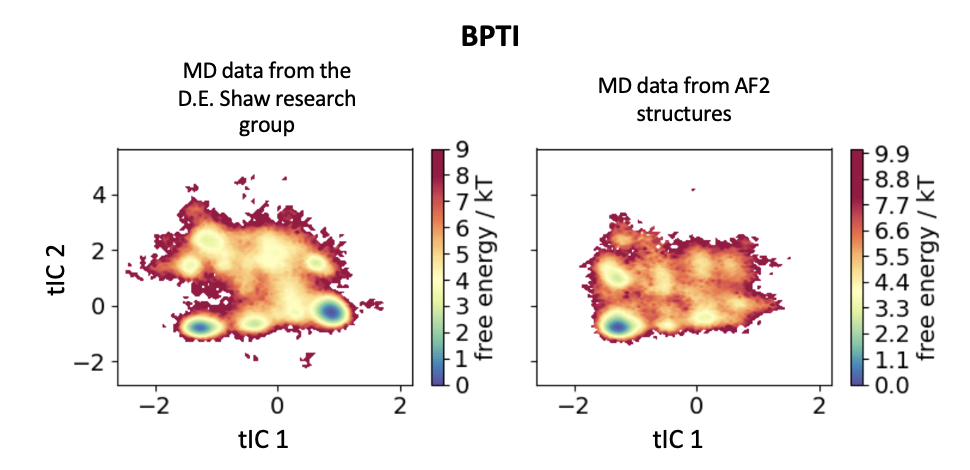

*Figure 3: Comparison of reweighted D.E Shaw MD data with reweighted seeded cMDs from the 32:64 without recycling subsampled ensemble of BPTI. The tICA is in a combined space of both simulations enabling the comparison of the represented free energy landscape and the corresponding minima.*

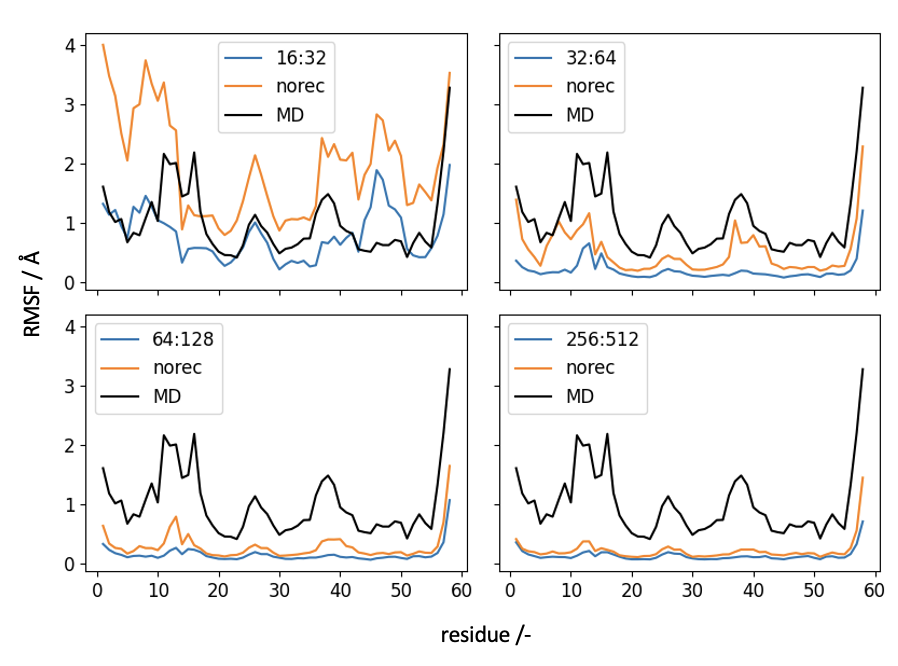

*Figure 4: Comparison of the residue-wise RMSF for BPTI of the AF2 ensemble predictions using different MSA subsamplings with (blue) and without recycling rounds(orange) to the MD simulations (black).*

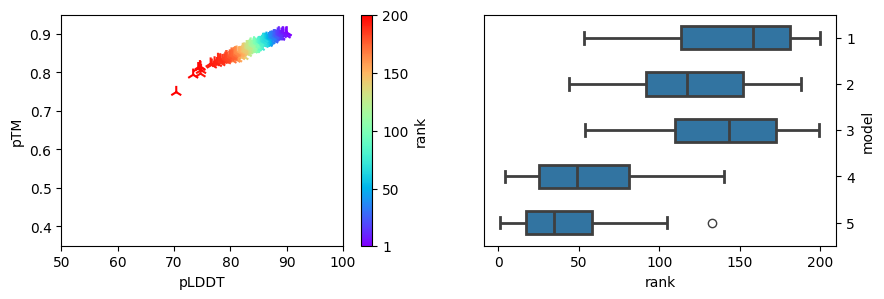

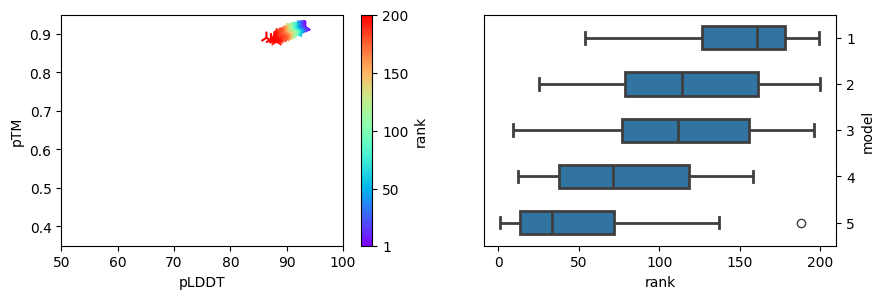

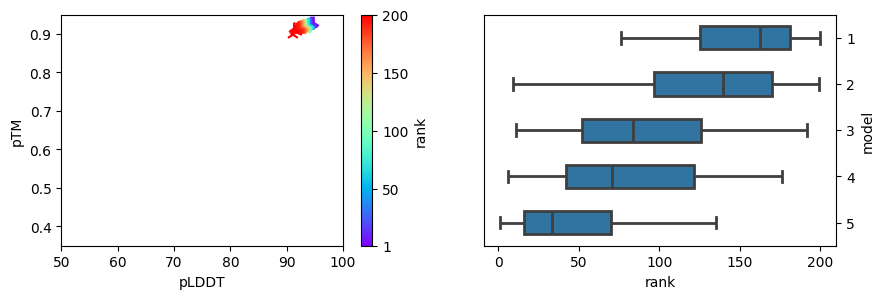

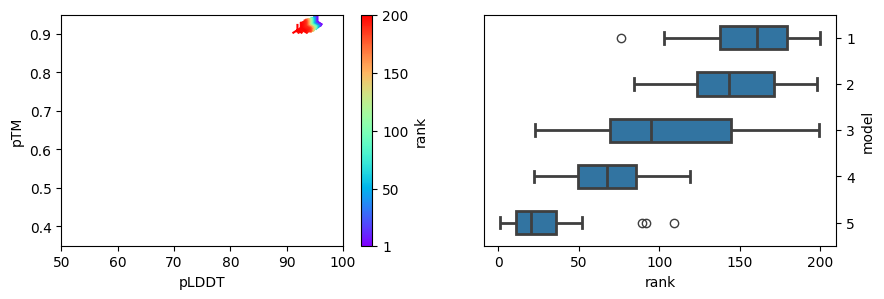

**16:32**

**32:64**

**64:128**

**256:512**

*Figure 5: Evaluation of the AF2 ensemble prediction of thrombin. On the left are the pLDDT and pTM score based on the rank depicted, whereas the right plots show the ranks of the different models.*

*Table 3: PDB accession codes for the selected thrombin structures in the active, zymogen and inactive form.*

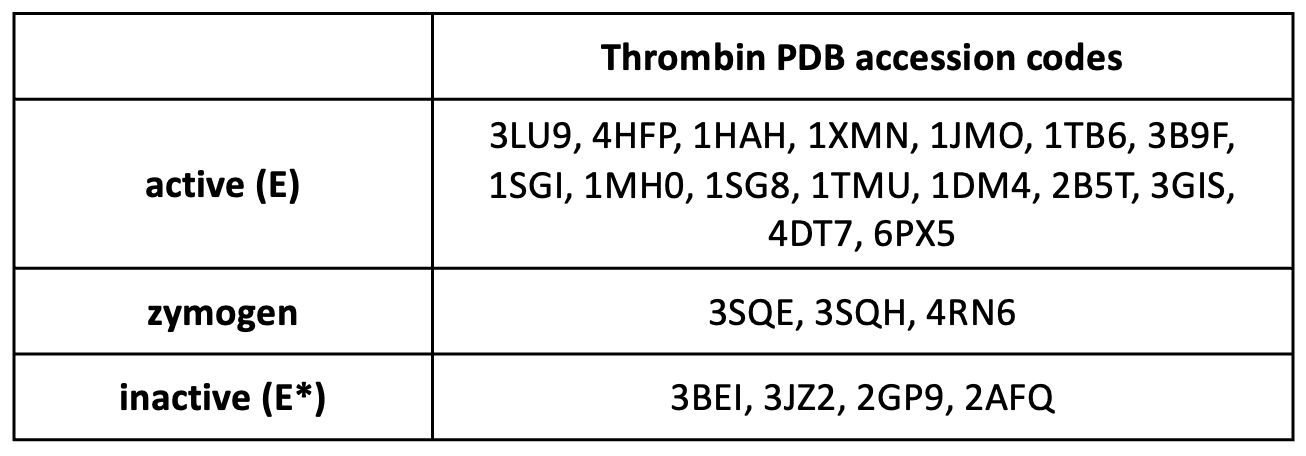

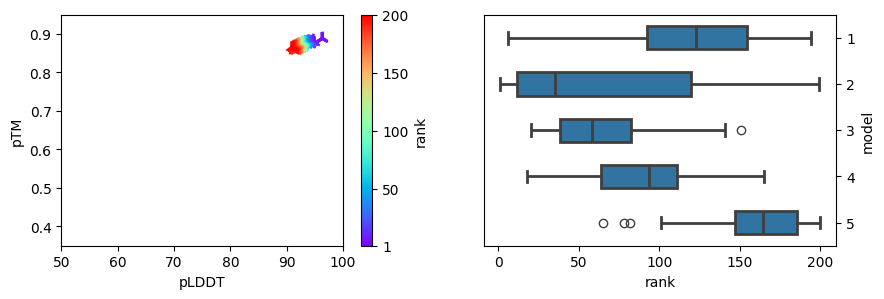

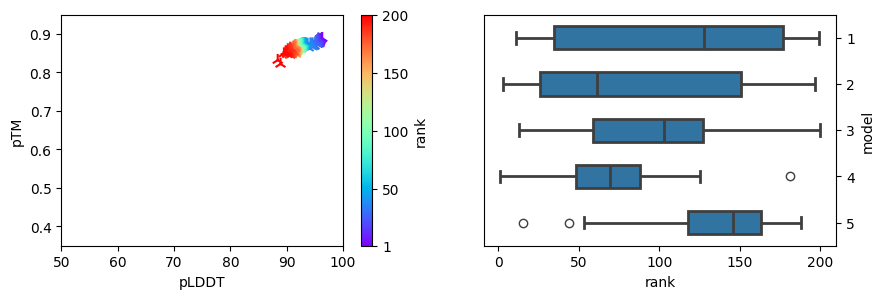

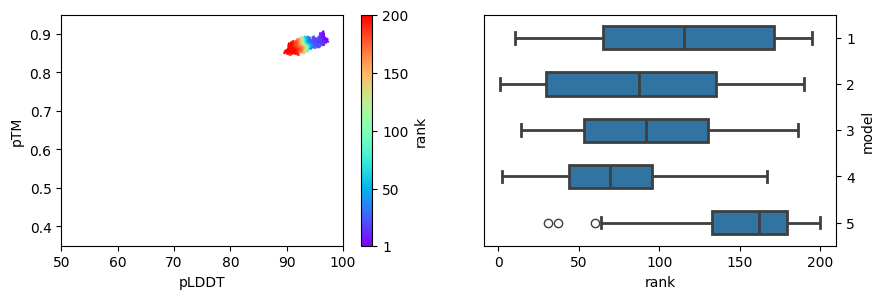

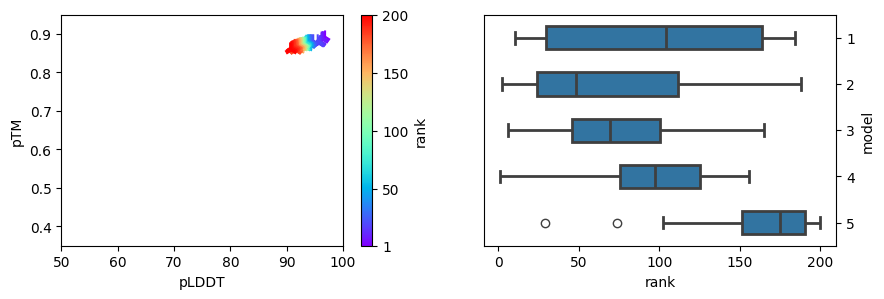

**16:32**

**32:64**

**64:128**

**256:512**

*Figure 6: Evaluation of the AF2 ensemble prediction of the nanobody. On the left are the pLDDT and pTM score based on the rank depicted, whereas the right plots show the ranks of the different models.*

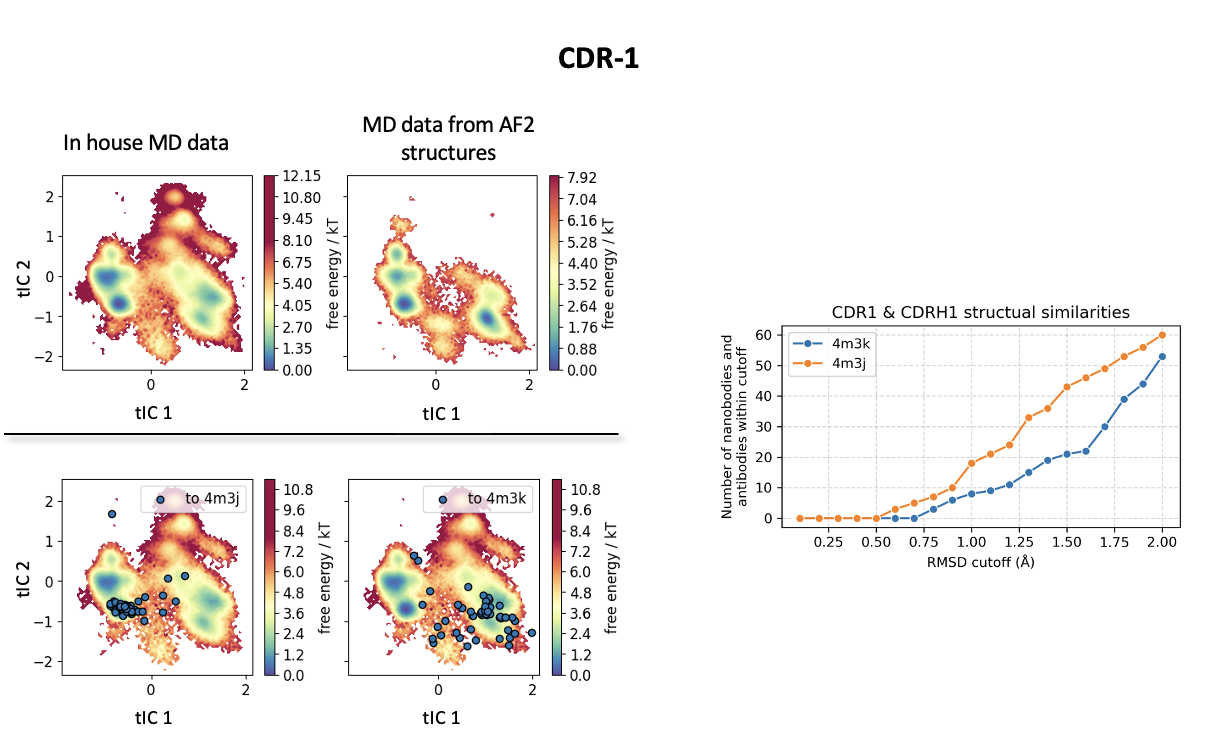

*Figure 7: Comparison of reweighted in-house MD data of CDR-1 with reweighted seeded cMDs from the 32:64 subsampled ensemble of the nanobody. The tICA is in a combined space of both simulations enabling the comparison of the represented free energy landscape and the corresponding minima. Additionally, we projected structurally similar structures to the crystal structures (PDB: 4M3K and 4M3J) present in the PDB (cutoff of up to 2 Å, in the right plot) in the tICA space of the in-house MD data (lower left).*

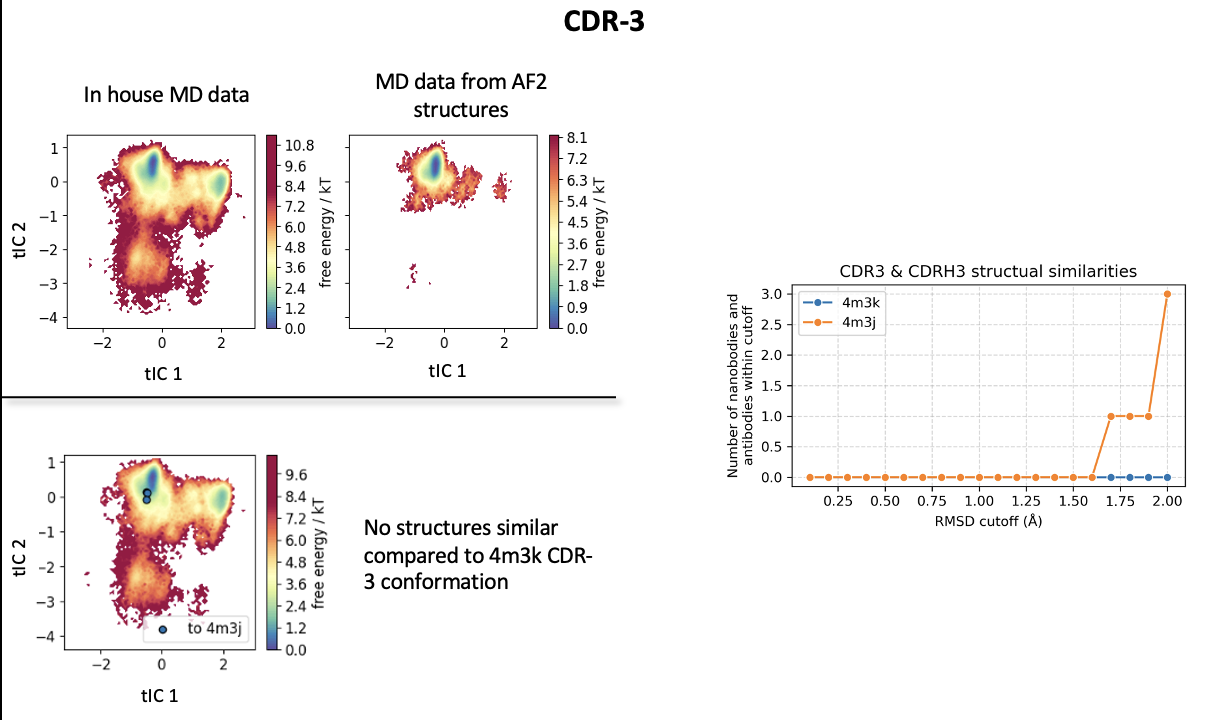

*Figure 8: Comparison of reweighted in-house MD data of CDR-3 with reweighted seeded cMDs from the 32:64 subsampled ensemble of the nanobody. The tICA is in a combined space of both simulations enabling the comparison of the represented free energy landscape and the corresponding minima. Additionally, we projected structurally similar structures to the crystal structures (PDB: 4M3K and 4M3J) present in the PDB (cutoff of up to 2 Å, in the right plot) in the tICA space of the in-house MD data (lower left).*

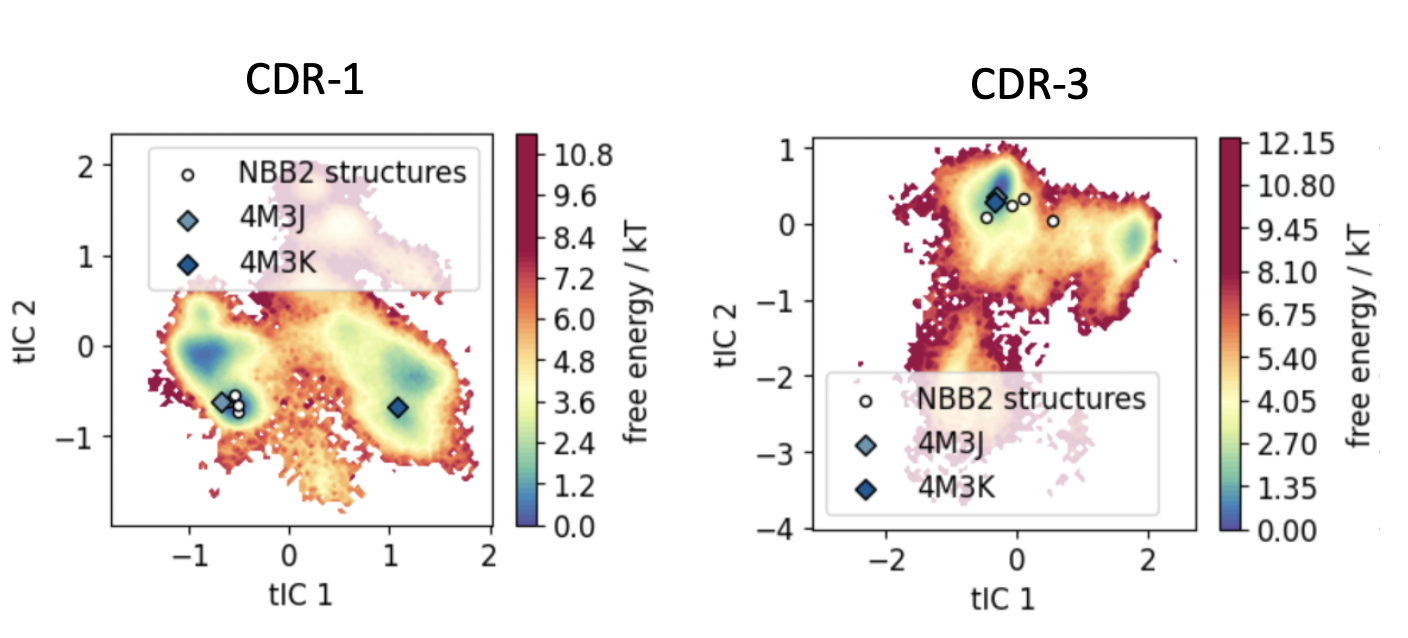

*Figure 9: Comparison of reweighted in-house MD data of CDR-1 and CDR-3 with the top four ranks from NanoBodyBuilder2 structure prediction (marked as white circles). The crystal structures (PDB: 4M3K and 4M3J) present in the PDB are represented as diamonds.*

*Figure 10: Evaluation of the AF2 ensemble prediction of the Fv. On the left are the pLDDT and pTM score based on the rank depicted, whereas the right plots show the ranks of the different models.*

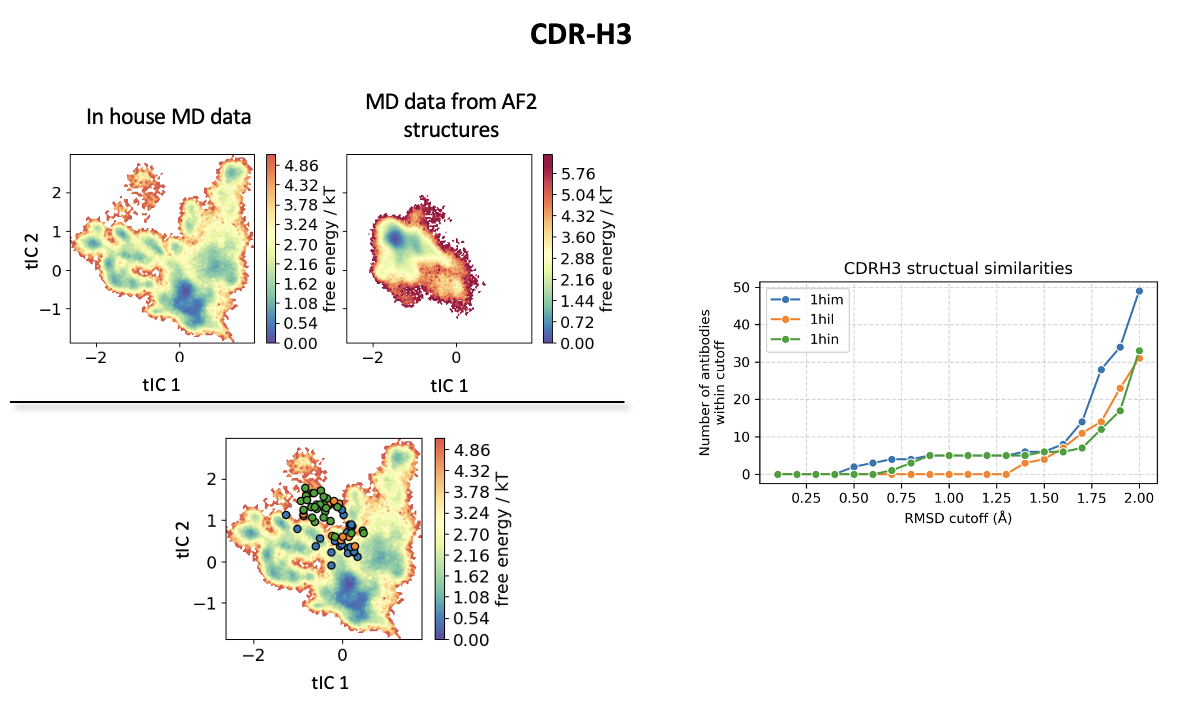

*Figure 11: Comparison of reweighted in-house MD data of CDR-H3 with reweighted seeded cMDs from the 32:64 subsampled ensemble of the Fv. The tICA is in a combined space of both simulations enabling the comparison of the represented free energy landscape and the corresponding minima. Additionally, we projected structurally similar structures to the crystal structures (PDB: 1HIM, 1HIL and 1HIN) present in the PDB (cutoff of up to 2 Å, in the right plot) in the tICA space of the in-house MD data (lower left, dots for the similar PDB structures are according to the right plot).*

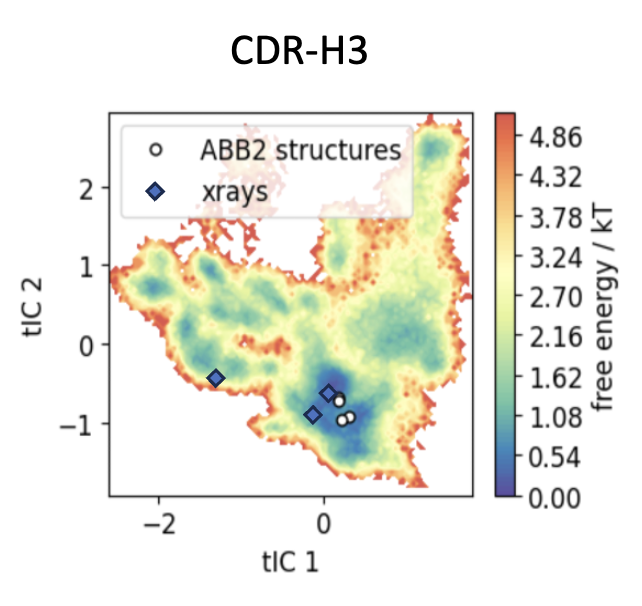

*Figure 12: Comparison of reweighted in-house MD data of CDR-H3 with the top four ranks from ABodyBuilder2 structure prediction (marked as white circles). The crystal structures (PDB: 1HIN, 1HIL and 1HIM) present in the PDB are represented as diamonds.*

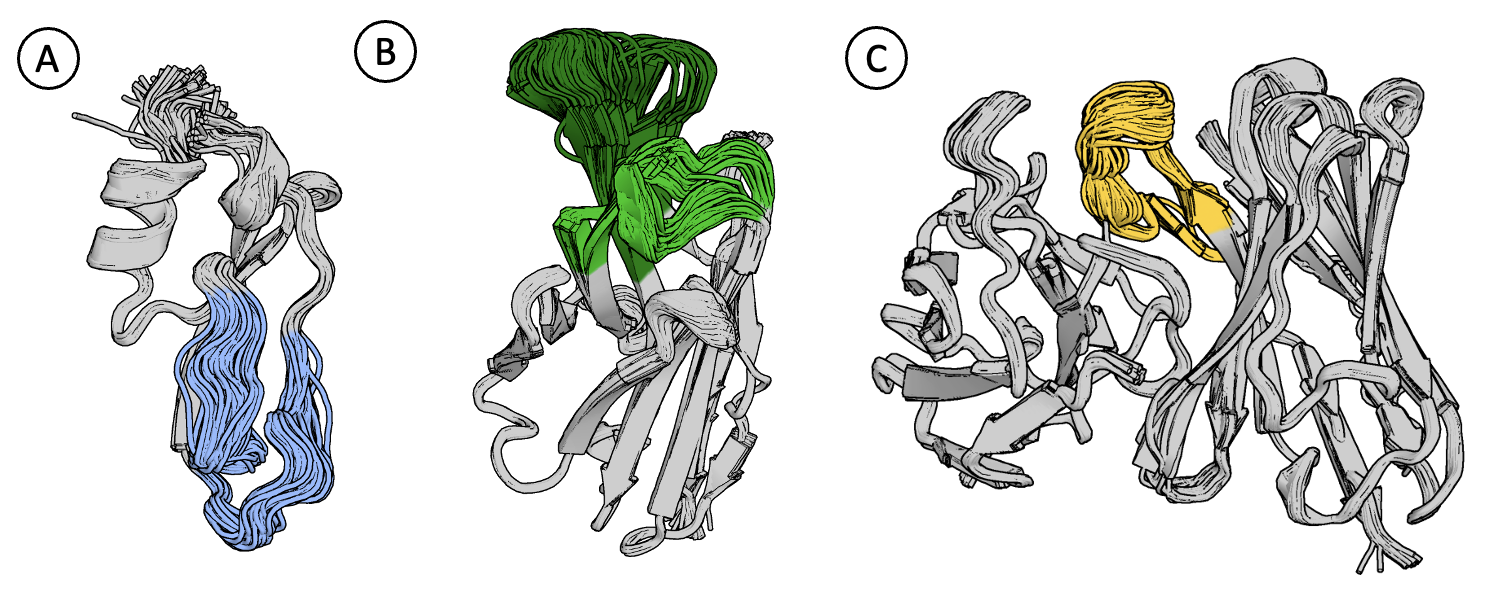

*Figure 13: Illustration of ensemble predictions that were further used as starting points for MD simulations. (A) represents the resulting ensemble from the 32:64 MSA subsampling without recycling of BPTI. The region around the disulfide bond isomerization is colored in blue. (B) represents the resulting ensemble from the 32:64 MSA subsampling of the nanobody. The CDR loops are colored in green. (C) represents the resulting ensemble from the 32:64 MSA subsampling of the Fv. The CDR-H3 is colored in yellow.*
